# Essentiality of local topology and regulation in kinetic metabolic modeling

**DOI:** 10.1101/806703

**Authors:** Gaoyang Li, Wei Du, Huansheng Cao

## Abstract

Genome-scale metabolic networks (GSMs) are mathematic representation of a set of stoichiometrically balanced reactions. However, such static GSMs do not reflect or incorporate functional organization of genes and their dynamic regulation (e.g., operons and regulons). Specifically, there are numerous topologically coupled local reactions through which fluxes are coordinated; and downstream metabolites often dynamically regulate the gene expression of their reactions via feedback. Here, we present a method which reconstructs GSMs with locally coupled reactions and transcriptional regulation of metabolism by key metabolites. The proposed method has outstanding performance in phenotype prediction of wild-type and mutants in *Escherichia coli* (*E. coli*), *Saccharomyces cerevisiae* (*S. cerevisiae*) and *Bacillus subtilis* (*B. subtilis*) growing in various conditions, outperforming existing methods. The predicted growth rate and metabolic fluxes are highly correlated with those experimentally measured. More importantly, our method can also explain the observed growth rates by capturing the ‘real’ (experimentally measured) changes in flux between the wild-types and mutants. Overall, by identifying and incorporating locally organized and regulated functional modules into GSMs, Decrem achieves accurate predictions of phenotypes and has broad applications in bioengineering, synthetic biology and microbial pathology.

---

Cellular life maintains itself and replicates through the entire set of biochemical reactions in genome-scale metabolic networks (GSMs) operating in a well-coordinated manner ^1-3^. But, how these reactions operate and are orchestrated and regulated *in vivo* in a GSM remains a major challenge in systems biology. Some insight into this question has been gained: local metabolite levels are stable in the face of environmental or genetic perturbations ^4-10^ due to quick compensation from nearby reactions. This suggests that rerouting of fluxes in GSMs operates efficiently, which plays a key role in metabolite homeostasis in maintaining global optima under one condition or across conditions ^5^. Furthermore, during the evolution of metabolic networks, reactions and metabolites prefer to actively attach to high-efficiency biochemical reaction chains in an organism, according to the principle of preferential attachment ^11-13^. This mode of network organization leads to topological coupling for metabolic flux rerouting. For example, in the TCA (tricarboxylic acid) cycle, the reaction chain D-isocitrate --> a-ketoglutarate --> … --> malate (*K*_m_=0.029μM and *K*_cat_=106.4 for IDH3 (Isocitrate dehydrogenase)) is preferred as the primary branch for oxidation of acetyl-CoA over the low-efficiency reaction chain D-isocitrate --> succinate (*K*_m_ =8μM and *K*_cat_ =28.5 for AceA (Isocitrate lyase)). Recent studies have revealed topology-derived module-specific enzyme parameters, e.g., *K*_m_ and *K*_cat_, network properties, e.g., path length and robustness, and gene expression profiles in GSMs ^14^. It has been demonstrated that some reactions are more closely interconnected with high kinetic capabilities than others and form small worlds in GSMs, which is the fundamental organizing principle of pathways ^15-17^, particularly in central metabolism, for essential precursor conversion in diverse environments ^18^. Therefore, coupled reactions constitute the topological foundation of local metabolic flux coordination in GSMs. The importance of these local coupling preferences is manifested in three aspects. First, the changes due to local internal perturbations (gene deletions) are quickly compensated by neighboring reactions ^10^. Second, functionally related genes are organized as operons that are coregulated in microorganisms ^19-23^. Third, failure to recognize the coupled reactions leads to poor performance in flux estimation by steady-state linear flux optimization, e.g., flux balance analysis (FBA) ^24^.

In response to internal (genetics) and external perturbations, metabolic systems are rather dynamic in adjusting metabolic fluxes ^5,25^, which results from regulations at the substrate and transcriptional levels ^26^. The mechanism of fast-acting and local substrate-level regulation of enzyme activity is well established, including allosteric regulation, feedback inhibition, and covalent modification ^27-29^. However, regulator-regulatee pairs of metabolites and enzymes are not available on the genome scale. Similarly, transcriptional regulation of metabolism on a genome scale has just begun ^8,30-33^. The expression profiles of metabolic genes are primarily regulated by the growth state via the sequestration or release of transcriptional factors (TFs) by key metabolites with concentrations that vary with growth ^32^. The activities of over 200 TFs show strong correlations with cognate metabolites following the transition from starvation to growth in *E. coli* ^*8*^. Compared to the global transcriptional regulation of metabolism, regulatory metabolites have not been identified on a genome scale, particularly in the functionally organized local modules of GSMs^8^. The regulation at the metabolic pathway level is particularly important because genes in microbial genomes are organized into operons (coregulated/coexpressed, functionally related and adjacently placed genes), and the expression of operons is regulated mainly by local TFs, which are in turn controlled by the end products of the pathways involving the genes of the operons ^8^. These facts underscore the potential of key metabolites directly controlling the pathway fluxes rather than simply being indicators of cell state with precise modeling of the associated enzyme (reaction) kinetics. Therefore, key metabolites may provide an alternative approach for reducing the complexity of dynamic flux prediction without loss in modeling accuracy, given the extreme difficulty of obtaining complete enzyme kinetic parameters (i.e., *K*_m_ and *K*_cat_) from paired metabonomics and proteomics data in building a reaction kinetics-constrained metabolic flux prediction model in GSMs.

In this study, we presented a computational method to integrate flux coordination derived from local coupled reaction topology and key regulator metabolite-mediated dynamic enzyme activity into GSMs to approximate biologically feasible fluxes (Fig. 1). We first incorporated the identified cooperated topological profiles of GSMs into the canonical FBA by representing the synchronously coordinated (coregulated) and closely coupled reactions with a group of independent sparse bases (reactions) according to the stoichiometric matrix decomposition. Then, we reconstructed a decoupled independent basis (reaction)-constrained metabolic model: Decrem, which was carried out on three model organisms: *E. coli, S. cerevisiae* and *B. subtilis*. The flux distributions predicted by Decrem were highly consistent with experimentally measured fluxes in various cellular behaviors across multiple strains: growth rate and the flux distribution on different carbon sources and in response to genetic and environmental perturbations. Next, a cell state-associated kinetic model (k-Decrem) was utilized to enhance Decrem by incorporating metabolite-derived TF regulation in response to environmental perturbation. A unique advantage of k-Decrem model is that pathway-associated key metabolite concentrations will suffice to achieve metabolic regulations and thus reduce the requirements for necessary kinetic parameters and paired multi-omics data. In addition, the accurate growth rates predicted by k-Decrem in *E*. coli genome-scale knockout strains revealed that intracellular perturbations are mainly ‘buffered’ by highly coupled reactions, which reveals the coordination between key precursors of central metabolism and cell growth. To the best of our knowledge, this is the first computational platform to integrate metabolic network topology and regulation by metabolites into GSMs.

**Fig. 1.**
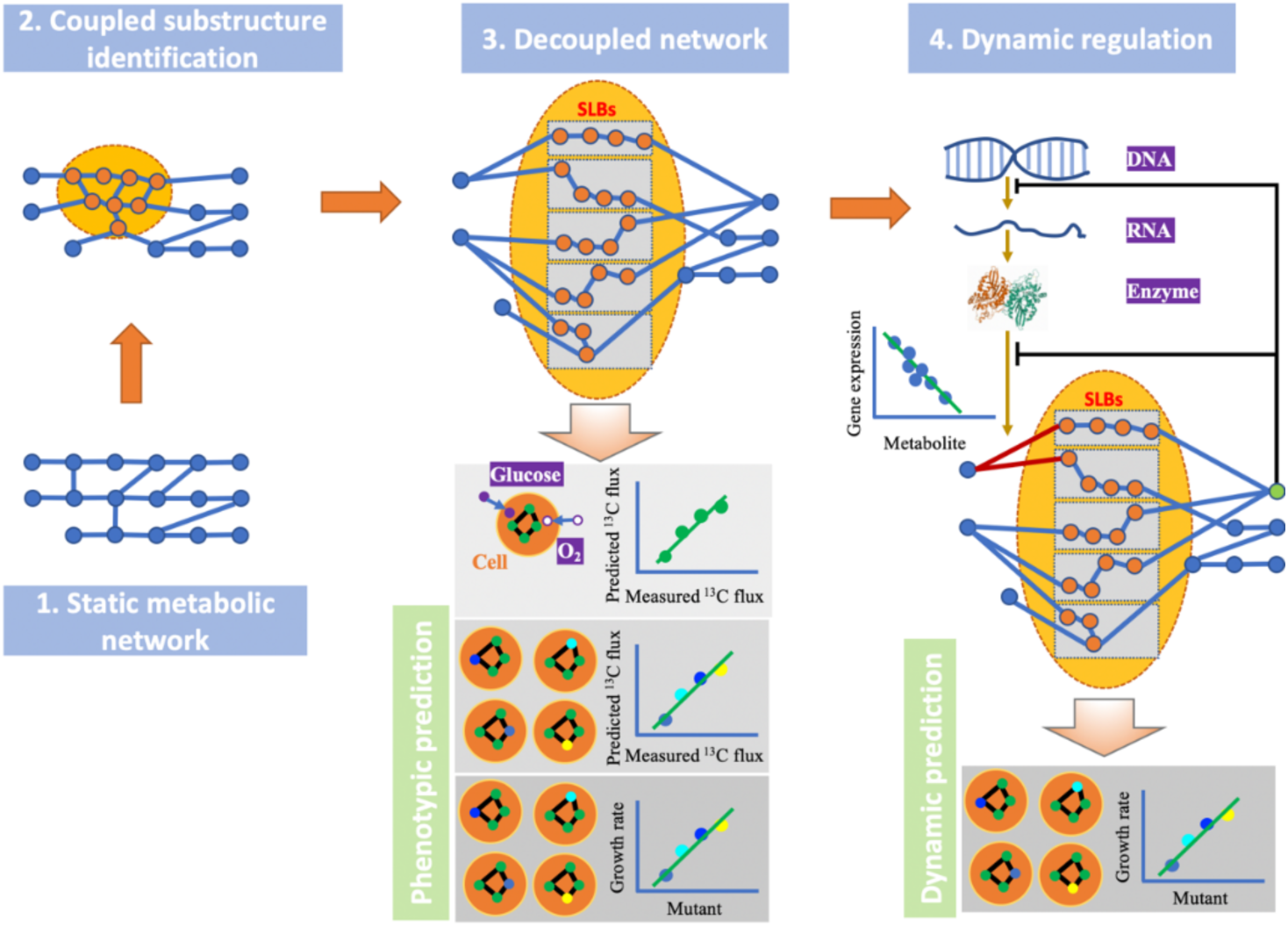
An overview of Decrem. (1) Schematic representation of a metabolic network. (2) Identification of a closely coupled reaction substructure (orange circles) of a metabolic network. (3) Closely coupled substructure is represented as a group of decoupled sparse linear basis (SLB) reactions (in gray rectangles), and the reconstructed GSMs incorporating these SLB reactions were tested in three scenarios: response in flux and growth rate to genetic perturbation and environmental perturbations. (4) Building a kinetic model integrating regulations at substrate and transcriptional levels, which is tested in growth rate prediction and is interpretation in genome-scale mutant strains.

## Results

### Reconstruction of GSMs with topologically decoupled reactions

We developed a topologically decoupled linear representation of metabolic network to characterize the coactivated regulation of highly coupled reactions with three steps. First, to explore the topological organization of highly coupled reactions, substructures composed of tightly connected coupled reactions in the metabolic network were identified from its bipartite graph representation with a graph clustering algorithm (see Methods; Supplementary Fig. 1) ^34^. Specifically, a similarity metric for coupled reaction substructure identification was defined as the number of simple cycles between any two reaction nodes in the bipartite graph. Then, reactions involved in multiple simple cycles were treated as candidate highly coupled elements and were classified into one of several coupled reaction-derived substructures using the minimum cut algorithm to further explore their linear decoupled representation (see Methods).

Specifically, identified coupled reaction substructures totally included 927 of the 2382 reactions in *i*AF1260 model of *E. coli* (Supplementary Table 1). Central metabolism, such as the TCA cycle, pentose phosphate pathway (PPP), glycolysis, and amino acid and glycerophospholipid pathways, primarily consists of coupled reactions and includes 70% (141 of 201) reversible reactions. In contrast, tRNA, membrane lipid biosynthesis, membrane transport pathways and the biomass reaction primarily consist of uncoupled linear reaction chains (Fig. 2a). Additionally, the *K*_m_ values (0.023 mM) of the coupled reactions in the substructures are smaller (by 56.5%) than those (0.036 mM) in the uncoupled reaction chains (Wilcoxon test, *p* = 5.74E-4; Fig. 2b). Together, the physical proximity and similar kinetic enzyme parameters suggest that these highly coupled reactions prefer to interact with each other first (as top priority), especially in central metabolism, before reaching out to reactions that are more distant. We then validated the coordination of the identified coupled reactions with publicly available fluxes experimentally measured in 24 knockout strains of *E. coli* ^35^. As expected, the correlations among the fluxes of the coupled reactions from the same pathways were higher than those for uncoupled reactions. For example, the average correlation coefficient *r* is 0.913, 0.975, and 0.794 for the reactions in glycolysis, PPP and TCA cycle, respectively, in contrast to 0.505, 0.267, and 0.421 for each uncoupled reaction set (*t* test, *p* = 2.33E-9, 3.06E-43 and 2.25E-4, respectively; Fig. 2c), which suggests that the topologically coupled reactions indeed also tend to coordinate their fluxes.

**Fig. 2.**
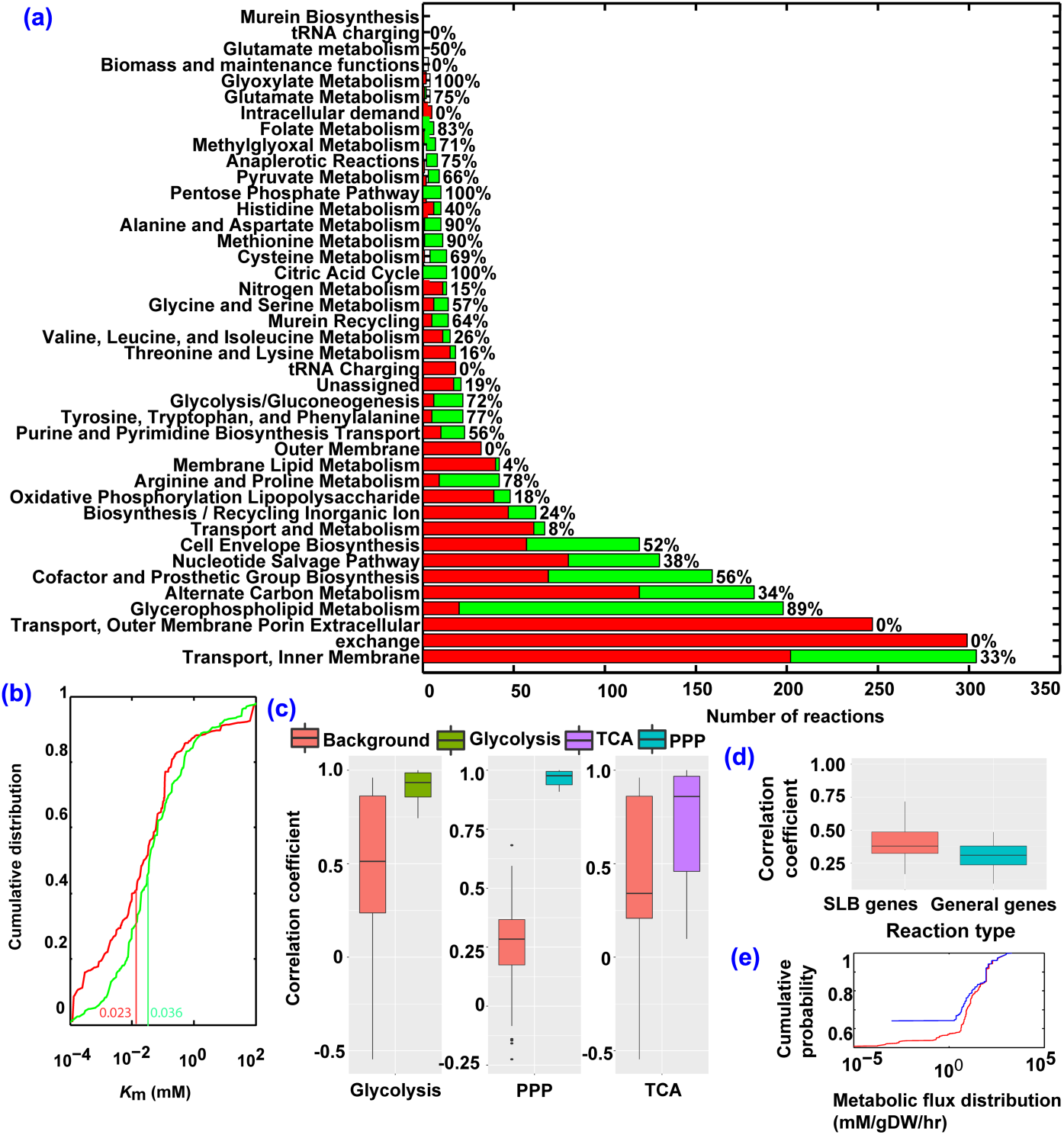
The statistical analysis of decomposed topological components of *E*. coli *i*AF1260 model. (a) The distribution of proportion of coupled reactions (green bar) among the 41 metabolic pathways. (b) The comparison of accumulated distribution of kinetic constant, K_m_, between the coupled reactions (red line) and non-coupled reactions (green line), the vertical bars are the mean K_m_ values of the two groups. (c) The correlation distribution of 13C MFA fluxes within the each of three pathways of central metabolism, compared to the corresponding distantly-coupled reactions (examined reactions from different pathways) in 24 strains. (d) The correlation distribution of gene expression among the element reactions of sparse linear basis (SLB) vectors against the gene expression among the non-coupled reactions; (e) The cumulative flux distribution of reconstructed decomposition metabolic model vs the original metabolic model according to flux variability analysis.

Second, we decomposed the highly coupled reaction substructures into representations with minimal independent components. Here, we represented the coupled reaction-derived substructures using sparse linear basis (SLB) vectors of the corresponding null space of the stoichiometry matrix and treated them as the decoupled components of GSMs. That is, each SLB consists of a least number of coupled reactions corresponding to an indivisible independent flux (see Methods). Like elementary flux modes ^36^, metabolic fluxes under real growth conditions could be decomposed as the weighted linear combination of the identified SLBs ^9^. However, unlike the almost infinite number of groups of ordinary linear bases, there is a unique group of SLBs in the null space of the stoichiometry matrix, which is an elegant way to define the mutually independent components for the densely interacting reactions ^35^ (Supplementary Information: Method S1). We validated the coordinated activation of reactions within the SLBs through gene expression in the 24 knockout strains of *E. coli* ^35^. Indeed, the correlations (mean *r* = 0.447) of gene levels among the element reactions from the same SLBs are higher than those (mean *r* = 0.28) from different SLBs from glycolysis, PPP and the TCA cycle (*t* test, *p =* 2.05E-33; Fig. 2d). This suggests that the reactions from the same SLBs tended to be coactivated, but the reactions in different SLBs are more independent.

In the last step, we reconstructed a model with decoupled linear representation of a GSM, which was named Decrem, by merging the element reactions of each SLB into a “linear basis reaction (LBR)” with reallocated stoichiometric coefficients (see Methods and Supplementary Fig. 1). To ensure that this decoupled representation has the same flux space as the original GSMs, flux variability analysis (FVA) was conducted in *E. coli* and *S. cerevisiae*. The results confirmed the preservation of solution space (Fig. 2e and Supplementary Fig. 2) ^37^. Our model reassigns the trivial fluxes while effectively preserving the primary fluxes of the original GSMs and serves as our working model for subsequent functional analyses.

### Prediction of metabolic fluxes in response to environmental changes

We first applied Decrem for metabolic flux prediction, in comparison with three other methods: FBA, pFBA (parsimonious FBA) and RELATCH ^38-40^ in three model organisms, namely, *E. coli* (*i*AF1260), *S. cerevisiae* (*i*MM904) and *B. subtilis* (*i*YO844), which are composed of 766, 558 and 332 LBRs in the corresponding decoupled GSMs, respectively (Supplementary Table 1, Data Availability).

We calculated the metabolic flux distribution in *E*.coli K12 MG1655 under four conditions in MOPS medium supplemented with glucose or xylose and under aerobic or anaerobic respiration ^41^ (Supplementary Information: Material S1), with nutrients as the sole constraints on GSMs (Supplementary Table 2). Using three evaluation metrics—Pearson correlation coefficient *r*, mean squared error (MSE) and the number of activated reactions, the predictions by Decrem have higher correlations with the experimentally measured ^13^C-MFA (metabolic flux analysis) fluxes (Supplementary Table 3) than the predictions by three other methods under all four conditions (Fig. 3). Decrem also produces much fewer discrepancies (and higher accuracy thereof) between predicted and empirical fluxes than the other methods (Fig. 3a). Furthermore, the fluxes predicted by Decrem have the smallest MSE (maximum upper bound of flux being 1000 mM/gDW/hr), and most of the activated reactions (with nonzero flux) agreed well with the experimental ^13^C-MFA fluxes (Fig. 3b and Fig.3c). Here, we provided an example of the superior performance of Decrem in the TCA cycle. The Decrem predictions are consistent with the experimentally measured ^13^C-MFA fluxes (Supplementary Fig. 3), whereas four activated reactions are predicted as inactive (zero fluxes) by FBA.

**Fig. 3.**
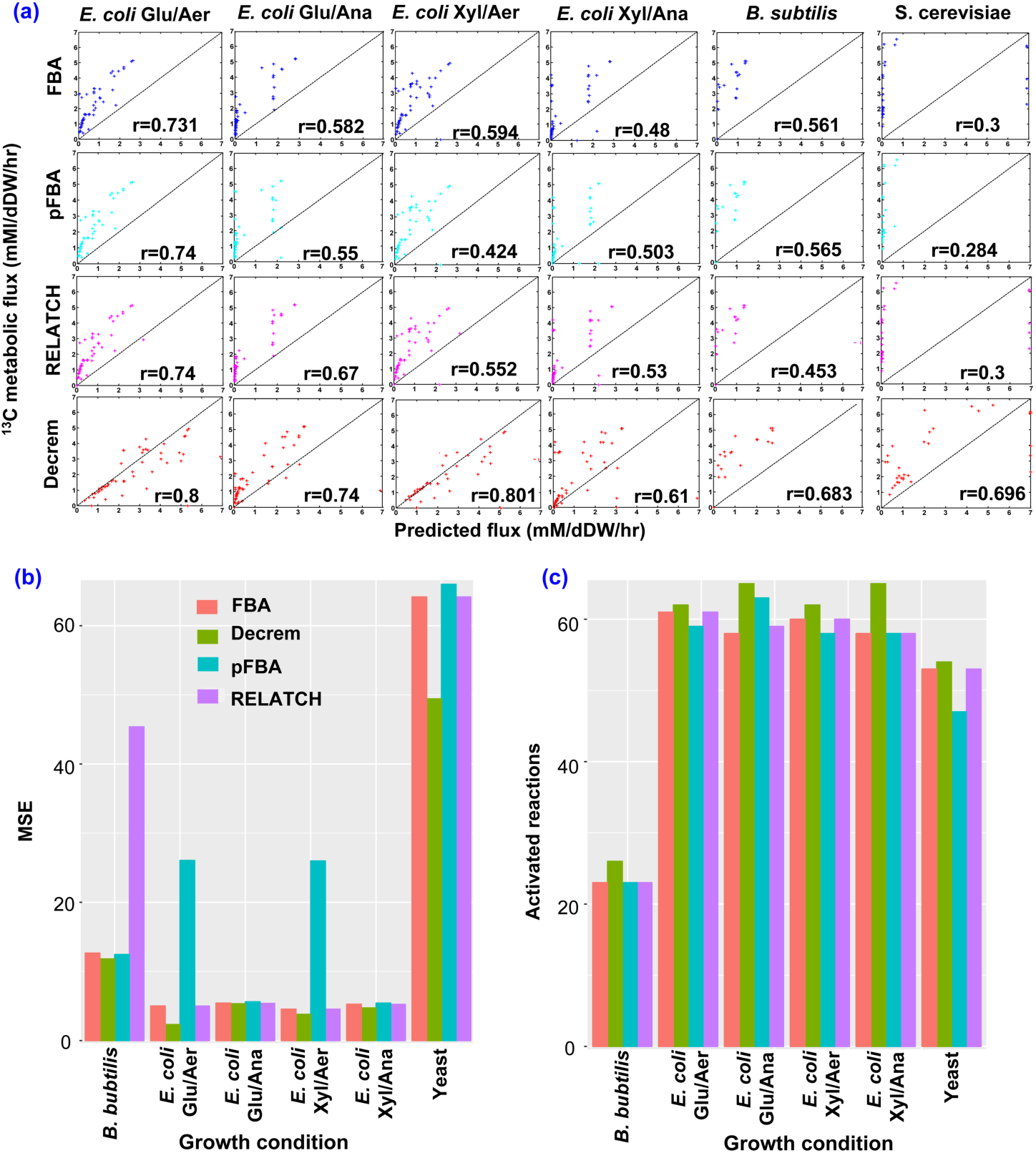
Comparison of the predictions in three model microorganisms by FBA, pFBA, RELATCH and Decrem. (a) Metabolic flux distribution between the experimental measurements and predictions by the four methods in *E*. coli *i*AF1260 with four conditions: glucose (Gly) or xylose (Oxy) under aerobic (Aer) or anaerobic (Ana) respiration, as well as in the *S*. cerevisiae *i*MM904 model and *B*. subtilis *i*YO844 model with glucose under aerobic respiration. r is the Pearson correlation coefficient between the measured ^13^C MFA fluxes and the predicted fluxes. (b) Comparison of MSE among the four methods across the six growth conditions. (c) Comparison of activated reactions among the four methods across the six growth conditions.

Decrem also outperforms the other three methods in flux prediction in two other organisms: *S. cerevisiae* and *B. subtilis* ^42,43^ (Fig 3; Supplementary Information: Material S1, Supplementary Table 1 and Supplementary Table 2). In particular, Decrem performs well with the complex eukaryote *S*. cerevisiae: *r* is 0.696, 0.3, 0.284 and 0.3 for Decrem, FBA, pFBA and RELATCH, respectively, in which reactions are often strongly coupled based on cellular compartments.

### Prediction of metabolic fluxes and growth rates in yeast knockout strains

In addition to fluxes in response to environmental changes, we further evaluated Decrem in predicting fluxes in response to genetic perturbation (single-gene deletions) in two GSMs of *S. cerevisiae, i*DN750 (1059 metabolites and 1266 reactions) and *i*MM904 (1226 metabolites and 1577 reactions). Thirty-eight single-gene-deletion mutants with experimental ^13^C-MFA fluxes, growth rates, nutrient uptake properties and several extracellular exchange fluxes ^44^ were used (Supplementary Information: Material S2; Supplementary Table 4). Decrem and four other methods: pFBA, FBA, RELATCH and REPPS ^45^ were used for flux prediction in these mutants. The predictions by Decrem showed the highest correlations with the experimentally measured fluxes in almost all mutant strains for both GSMs (Fig. 4 and Supplementary Table 5). Specifically, the average Spearman and Pearson correlation coefficients (*r*) determined by Decrem for *i*DN750 are 0.72 and 0.763 in the 38 mutant strains, compared to 0.615 and 0.733, 0.607 and 0.681, 0.556 and 0.707, 0.462 and 0.595 for pFBA, FBA, RELATCH and REPPS, respectively (one-way ANOVA, *p* = 5.52E-20 and 4.21E-07). For the more complete yeast *i*MM904, the mean Spearman and Pearson’s correlation coefficients are 0.782 and 0.936, 0.765 and 0.747, 0.656 and 0.915, 0.75 and 0.925, 0.645 and 0.841 for Decrem, pFBA, FBA, RELATCH and REPPS, respectively, which are significantly different (one-way ANOVA; *p* = 4.35E-06 and 8.26E-06, respectively). The differential accuracy of flux predictions between the *i*DN750 and *i*MM904 models shows that Decrem performed better than other methods, particularly on metabolic models that are less complete (e.g., *i*DN750 vs *i*MM904) (Fig 4). Further analysis revealed that such differences exist because a large proportion, 47% (146 of 311), of new reactions in *i*MM904 (vs *i*DN750) are highly coupled reactions, compared to an average 35% (558 of 1577) for *i*MM904 (Fisher’s exact test, *p* = 5.427e-04; Data Availability), which clearly optimizes the solution space of the metabolic model and changes the optimal reaction paths in *i*MM904.

**Fig. 4.**
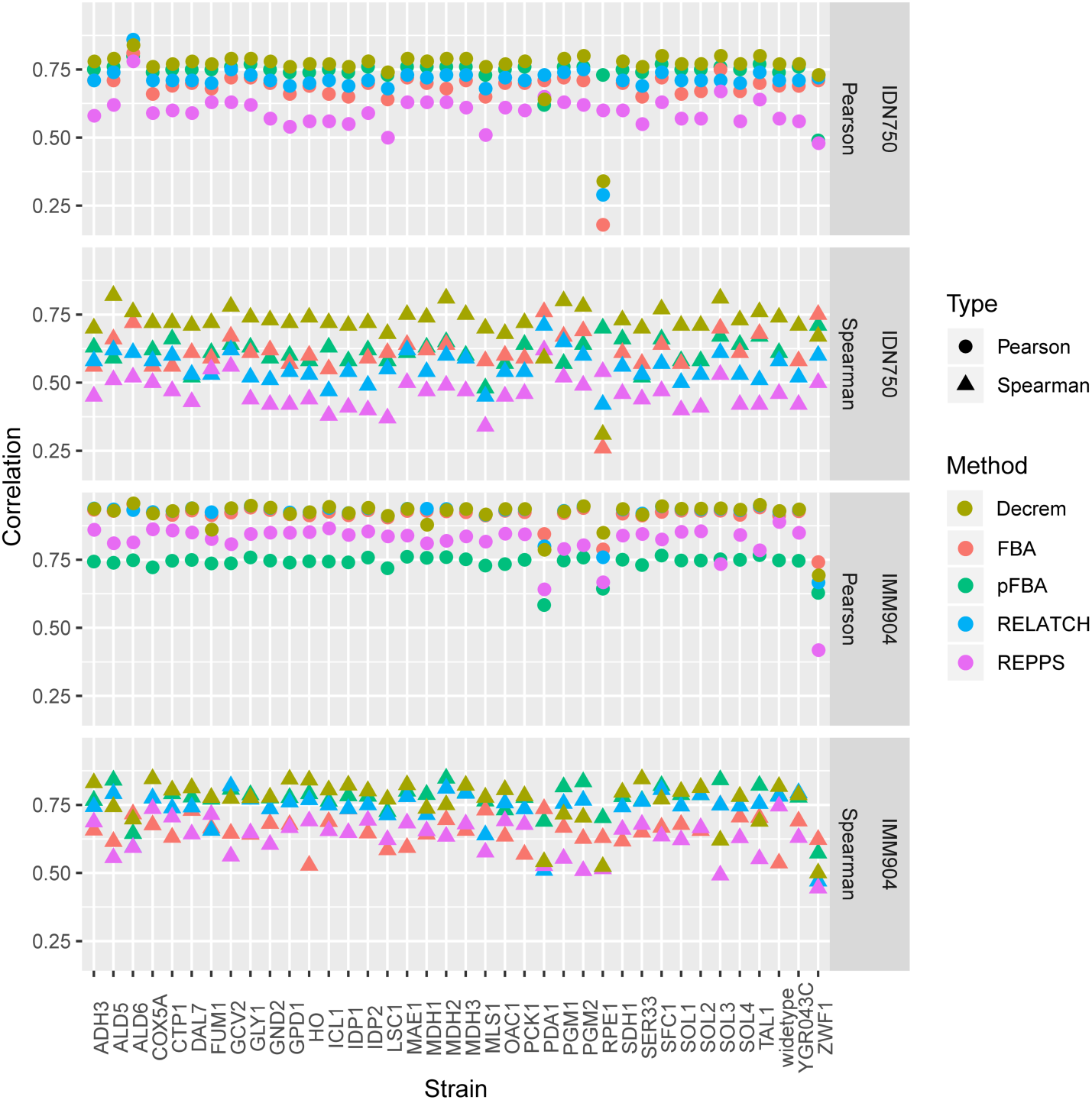
Comparison of the flux predictions in two *S. cerevisiae* metabolic models. The Pearson and Spearman correlation between the ^13^C MFA measured and predicted metabolic fluxes across each of the five test methods (FBA, pFBA, RELATCH, REPPS and Decrem) among 38 strains in the *i*MM904 model and *i*DN750 model of *S. cerevisiae*.

We then applied Decrem and pDecrem (parsimonious Decrem, see Supplementary Information: Method S2) to estimate the growth rate in the 38 mutant strains of *S. cerevisiae* using the *i*MM904 model. PCA of the predicted fluxes from all six checked methods shows that the top two principal components (PCs) can explain more than 99% of the variance of all predicted fluxes. The two PCs of Decrem predictions are highly correlated (*r* > 0.7) with experimentally measured growth rates, while the other methods have, at most, one PC that merely shows moderate correlation (*r* ∼ 0.5) (Supplementary Table 5). Furthermore, a PCA regression reveals that Decrem produces the best flux variance to explain the observed growth rates, as shown by the coefficient of determination (R^2^), which was 0.74, 0.731, 0.731, 0.841, 0.9 and 0.9 for the six methods (Fig 5a). Most importantly, Decrem successfully identified all the mutations with large effects leading to growth rate < 0.5 h^−1^: ALD6, FUM1, PDA1, RPE1 and ZWF1 (Fig 5b). Moreover, Decrem-predicted flux distribution through the specific pathways could explain the measured low growth rates. For example, the ZWF1 mutant had a different flux distribution from those of ALD6 and RPE1 (Fig 5c). Interestingly, the exceptionally high fluxes of the mitochondrial transport pathway and citric acid cycle pathway in the ZWF1 strain agree well with the experimental observation that NADPH- and NADP^+^-dependent mitochondrial malic enzyme flux is significantly increased ^44^. The other methods only identified some large-effect mutations on growth rates, leading to a high false-positive rate. In comparison, Decrem demonstrated high accuracy and low false positive rates in assessing the growth rate and well approximated the real metabolic fluxes in the mutants well.

**Fig. 5.**
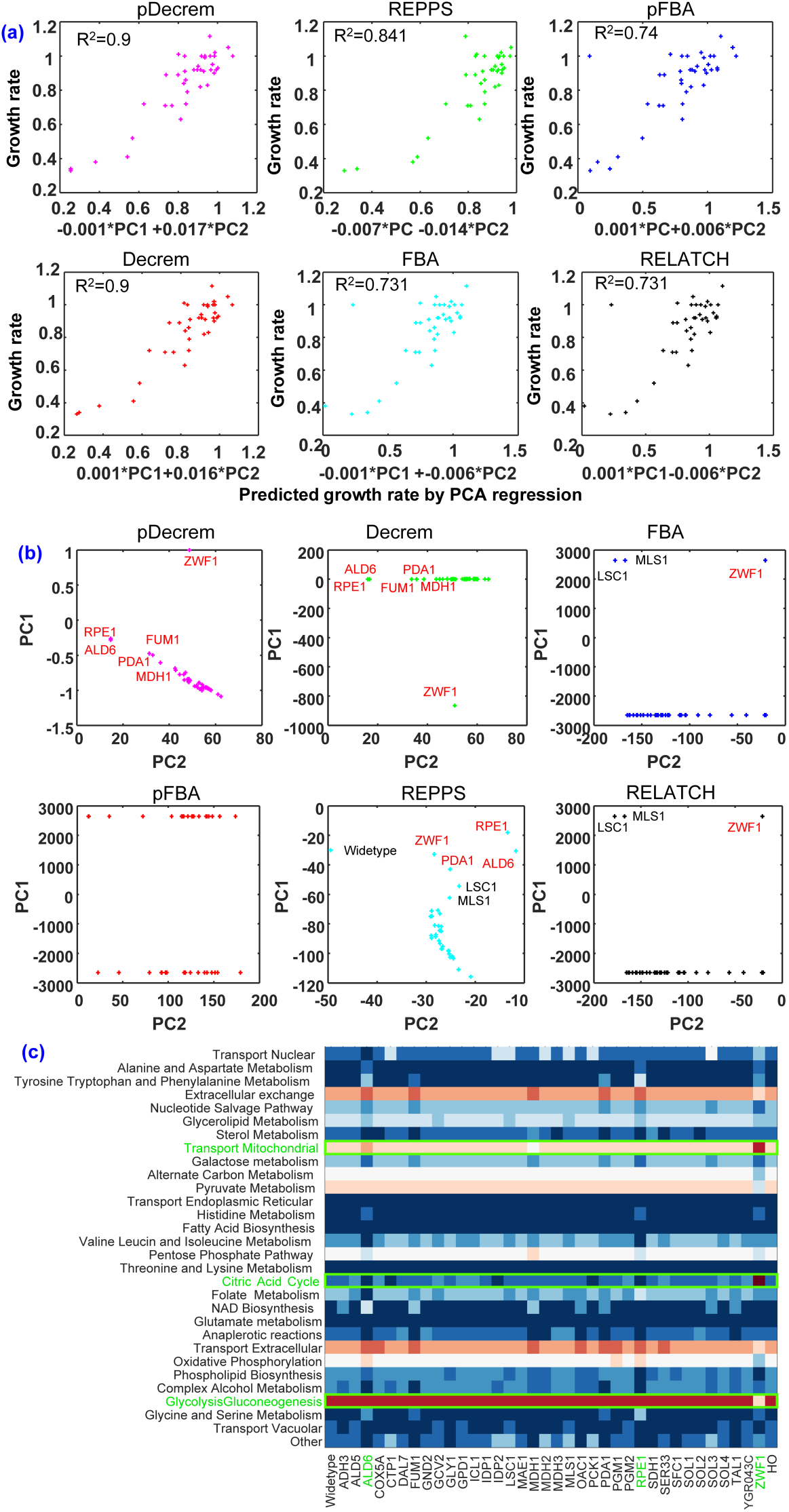
Cell state predictions for S. cerevisiae by FBA, pFBA, RELATCH, REPPS, Decrem and pDecrem. (a) representation of the experimentally measured (y-axis) and predicted (x-axis) growth rates with PCA regression of the top two PCs from the six test methods. (b) The top two principal components (PCs) of predicted flux variances among the 38 mutant strains, according to the six test methods. The deleted genes with high negative effects on growth are colored in red and the genes with no significant effects on growth are colored in black. (c) The heatmap of predicted fluxes by Decrem through 30 pathways revealed different responding mechanisms among the ZWF1 and the ALD6 mutant, as well as the RPE1 (highlighted pathways in green box indicates the ZWF1 deletion strain specific flux distribution).

### Building a metabolic kinetic model incorporating metabolite-TF regulation

Cellular metabolism is not static but rather dynamic because of regulation by substrates and transcription in response to internal and external perturbations. However, for transcriptional regulation of metabolism on a global scale, few regulator-regulatee pairs of metabolites and genes are known. Here, we identified the regulatory roles of key metabolites in central metabolism by incorporating highly coupled coactivated reactions into a metabolic kinetic GSM ^46-48^, which is verified with empirical multi-omics data in *E. coli* (see Supplementary Fig. 4).

First, we applied a quantitative approach to identifying pairs of regulator metabolites and regulated genes. As the TFs of metabolic genes in central carbon metabolism are regulated by global growth rate and local pathway-specific essential metabolites, e.g., AMP, fructose-1,6p, and fructose-1p in *E. coli* ^48^, we hypothesized that there are quantitative relationships between the expression levels of metabolic genes regulated by these TFs and the biomass constituents (growth rate being equivalent to change in biomass), as well as essential metabolites. We then tested this hypothesis with a multi-omics dataset of 24 single-gene knockout strains of *E. coli* ^35^, which includes the expression of 85 metabolic genes in central metabolism, concentrations of over 100 metabolites and 51 ^13^C-MFA fluxes for each strain ^49^. Through an extensive literature survey ^50,51^, 45 essential metabolites (and biomass constituents) were selected as potential regulators. They were classified into two groups according to their concentrations: the biomass-constituent group (BG) and the precursor or essential metabolite group (PG) ^2^ (Supplementary Fig. 5 and Supplementary Table 6). This classification displayed different concentration patterns between the global (biomass constituents) and local (pathway-specific) regulators (Supplementary Information: Material S3).

Next, we conducted a partial least squares regression (PLSR) to fit the observed expression profiles of the 85 genes to the concentrations of the potential regulatory metabolites of either group (see Methods). By taking stringent combined thresholds (total regression correlation *r* > 0.84 and the correlation of first PC > 0.38 according to PLSR; Supplementary Information: Material S3), 32 of the 85 genes were identified as the regulatory targets of the 23 BG metabolites, whereas no genes were identified as the regulatory targets of the PG metabolites (Fig. 6a and Supplementary Fig. 6). These identified metabolite-gene regulatory pairs are highly consistent with quantitative proteome-derived metabolite concentrations and the target genes of well-known metabolite-regulated TFs (Supplementary Table 6) ^48,52^. The identified metabolite-gene regulatory pairs are further verified with correlations between the measured and predicted gene levels using a wide range of metabolites selected by 10,000 random samplings from the 45 potential metabolic regulators (see Supplementary Information: Material S3). A *p* value of 3.1E-3 for the 23 identified growth-associated metabolites was observed against the randomly selected metabolites (Fig. 6b), while the *p* value was 0.48 for the PG metabolites.

**Fig. 6.**
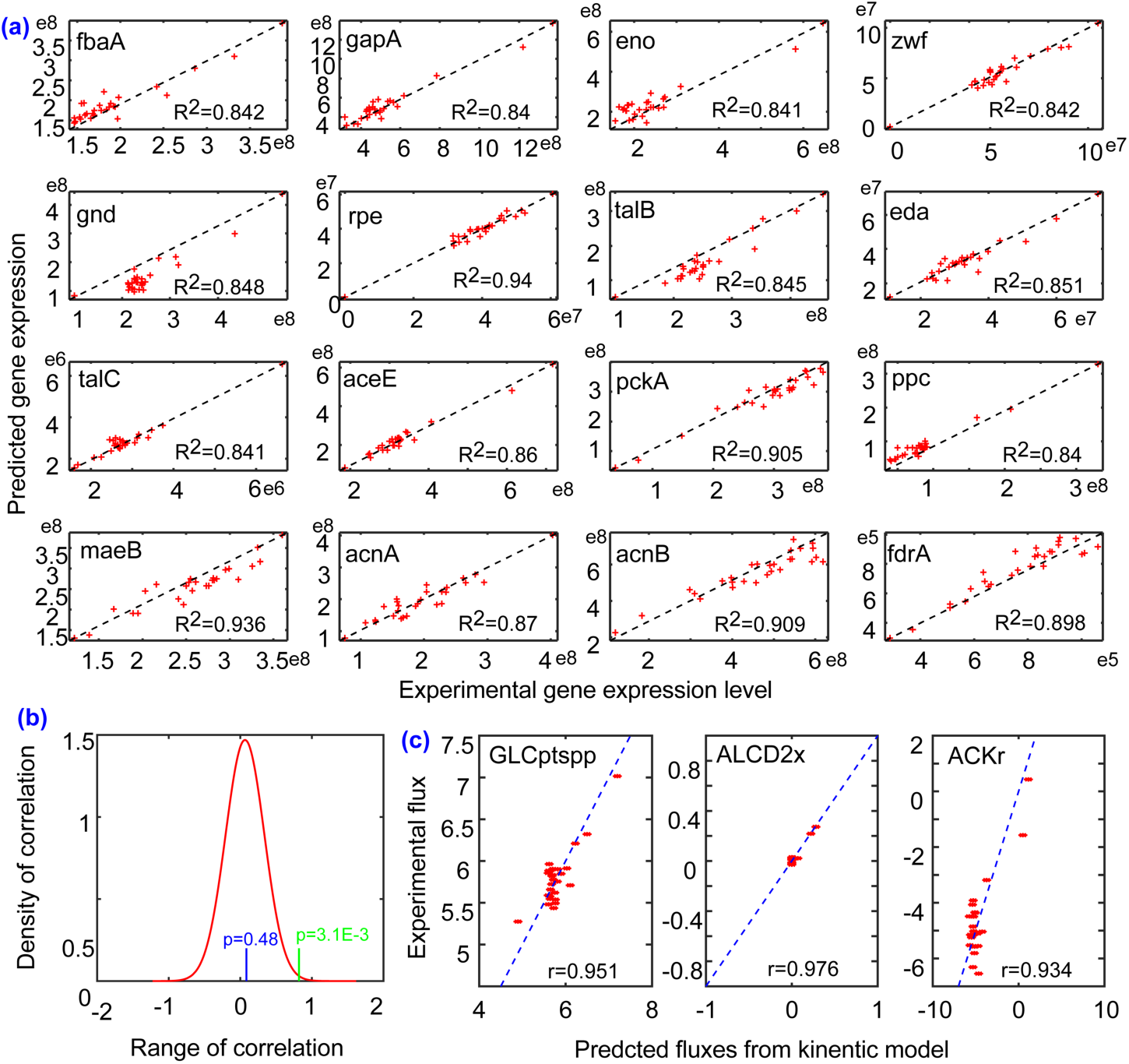
The analysis of metabolites-based kinetic regulation model of *E*.coli in 24 mutant strains. (a) The predictions of 16 genes expression using the 23 biomass-associated metabolites by the partial least squares regression. (b) The distribution of correlation between the measured and predicted gene expression levels using the metabolites from 10,000 random sampling (red curve), with the statistical significance levels according to the biomass metabolites group (green line) or the precursor metabolites group (blue line). (c) Predicted kinetic fluxes for three key regulatory reactions according to metabolites-determined Michaelis-Menten kinetics.

The identified 32 genes are primarily located in the PPP and pyruvate metabolism in KEGG pathways (Fig 7), which are associated with cell growth for biosynthesis: generating NADPH and pentoses toward nucleotide and amino acid biosynthesis ^53^, instead of being in energy-producing pathways (TCA and glycolysis). These results suggest that the expression of the genes in growth-associated pathways could be represented as a linear combination of the concentrations of biomass composition.

**Fig. 7.**
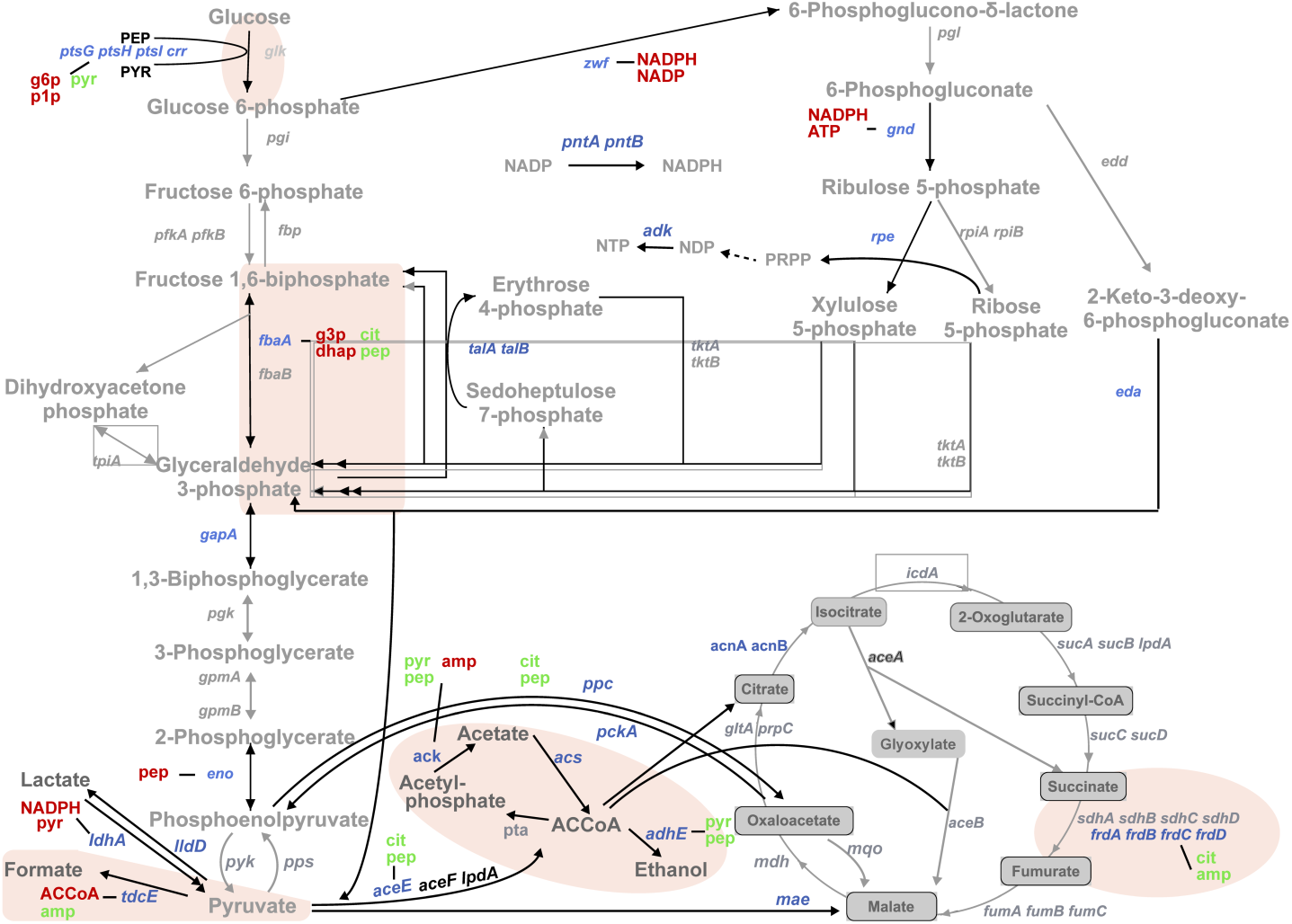
A map showing of regulator metabolites and the genes regulated by the biomass-constituent metabolites in the central metabolism. The meaning of different backgrounds and colors: red and green metabolites are gene inhibitors and activators; the blue genes present the identified 32 global growth state associated genes. The brown background indicates the identified key regulated reactions which are used to build the kinetic Decrem.

We then constructed a regulation-enabled kinetic model, k-Decrem, for the identified growth-associated metabolic reactions depend only on the concentration of corresponding metabolites above (see Methods; Supplementary Information: Material S3) and evaluated it by optimizing the growth-associated key regulated fluxes in *E. coli* central metabolism. The predicted optimal kinetic fluxes from the kinetic model display high consistency with experimental measurements through ^13^C isotope tracing (Fig 6c), with *r* values of 0.95, 0.97 and 0.93 for the reactions of glucose transport, alcohol dehydrogenase and acetate kinase, respectively, which are experimentally measured. These fluxes are also utilized to assign key kinetic fluxes in genome-scale metabolic flux prediction below (see Methods).

### Growth rate estimation for *E. coli* genome-scale gene deletion strains with k-Decrem

We applied k-Decrem constructed above to predict the growth rates of *E. coli* genome-scale single-gene deletion mutants, the growth rates and the corresponding concentrations of over 7,000 metabolites of which have been experimentally measured ^54^. A total of 1,030 mutants with genes involved in metabolism were selected for growth analysis (see Supplementary Information: Material S4). We first examined the growth rates predicted by the kinetics-free methods pFBA, MOMA and Decrem without any external flux constraints. As expected, poor results are produced, with low correlations with experimentally measured growth rates and *r* values of 0.127, 0.103 and 0.281 for pFBA, MOMA and Decrem, respectively (Supplementary Fig. 7).

Next, we constructed the kinetic fluxes of five key regulated reactions in central metabolism with predictions (based on metabolite concentrations) for each of 1,030 mutants to approximate the gene mutant-specific metabolic state, e.g., the branch points of glycolysis and PPP, the flux allocation downstream of pyruvate metabolism, and the growth-associated secretion (see Methods; Fig. 7 and Supplementary Table 7). On the GSMs integrated with kinetic fluxes, the growth rates of the 1,030 mutants were estimated with six methods: Decrem, pDecrem, FBA, pFBA, RELATCH and REPPS. The results showed that all six kinetic methods have significantly improved predictions compared to the kinetic-free methods (Fig 8a). Among them, k-Decrem and k-pDecrem produced the highest correlations with the empirical growth rates (*r* = 0.731 and 0.743 vs 0.421, 0.685, 0.474, and 0.509) (Supplementary Table 7).

**Fig. 8.**
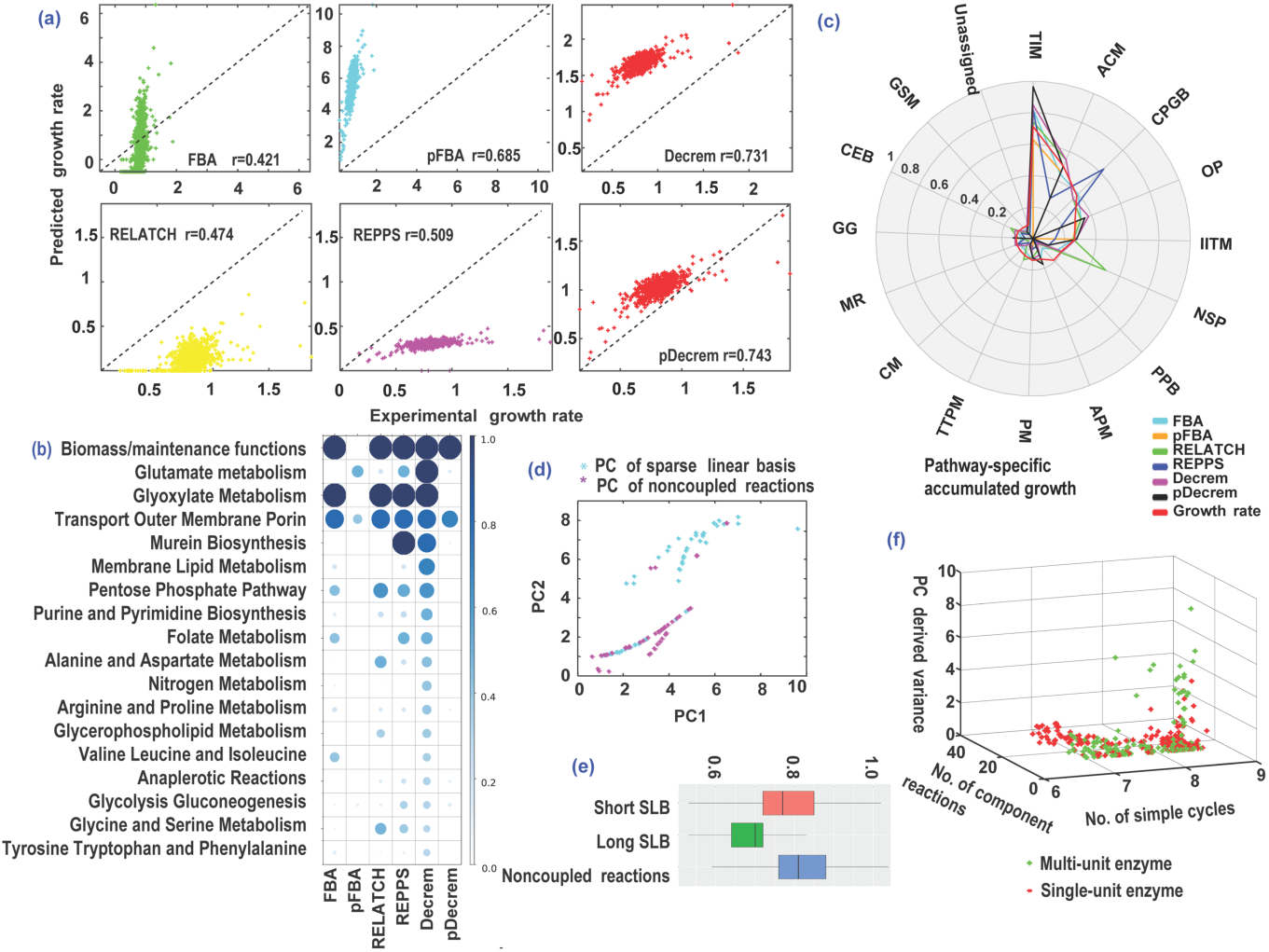
The growth rate analysis on the genome-scale gene deletion strains. (a) The comparison of measured and predicted growth rates by the k-pDecrem, Decrem, FBA, pFBA, RELATCH and REPPS, on the 1,030 mutants. (b) The correlation between the measured growth rates and the predicted accumulated fluxes though the identified growth-associated pathways. (c) The comparison of the predicted accumulated growth rates through the pathways where the mutants belong, according to the six test methods. (d) The top two PCs revealed difference in flux variances between the coupled reactions (cyan dots) and the non-coupled reactions (purple dots) which are predicted by the Decrem according to the 1,030 strains. (e) The comparison of growth rates distribution among topological components where the mutants belong: the complex LBR (the number of element reactions of corresponding sparse linear basis vector > 1), the simple LBR (the number of element reactions of corresponding sparse linear basis vector = 1) and the non-coupled reactions. (f) The flux variance of LBR based on the multimeric classification: the multi-subunit enzymes (green dots) and single-unit enzymes (red dots), the number of simple cycles and the length (the number of element reactions) of LBR (Fisher’s Exact Test, p values = 2.34E-21 and 0.0051).

We then demonstrated the explanatory power of k-Decrem in interpreting the observed growth rates of mutants with corresponding (altered) flux distributions. For that, we calculated the distribution of pathway-specific fluxes across mutant strains, defined as the accumulated flux (AF) ^9^ of each pathway (the accumulated sum of all nonzero fluxes in a pathway for each mutant strain) (Supplementary Information: Material S4). The correlations between the AFs and growth rate for all strains showed that k-Decrem quantified the greatest number of growth-related pathways that we curated from the literature compared to the other methods. For instance, many well-known important pathways for cell growth—glutamate, nucleic acid and most amino acid metabolism pathways—are ‘detected’ only by k-Decrem with large correlation coefficients (Fig 8b). Interestingly, although the kinetic pFBA (also k-pDecrem) method can predict the growth rates with relatively high accuracy, the corresponding AFs covered only a few of the curated growth-associated pathways. Moreover, the analyses of the pathway-specific accumulated growth rates (AG; the accumulated sum of growth rates of strains in which the mutated gene is located in the same pathway) suggest that k-Decrem predicted distributions of AGs through all metabolic pathways are highly consistent with the experimentally measured AG distributions (Fig 8c). FBA reaches similar levels of accuracy, but the predictions by pFBA only cover AGs, which were weakly influenced by gene knockouts and shrank the AGs to zero for the pathways containing knockout genes with strong effects (Fig 8c). Such biases are an intrinsic property of the L1 norm-based pFBA (and pDecrem) method ^54,55^, despite the relatively good fit of correlations.

We further examined the flux variance distribution in each mutant for their changed growth rates. PCA of the fluxes predicted by k-Decrem across the 1,030 mutants is shown in Fig 8d (Supplementary Information: Material S4). The distribution of the top two PCs indicates that the primary flux variances come from the decoupled LBRs, compared to the uncoupled reactions: 1.461 vs 1.10 on average for PC1 (*t* test, *p* = 3.04E-20); 1.56 vs 1.07 on average for PC2 (*t* test, *p* = 2.86E-25). This is consistent with the high robustness of the central metabolism ^14,56^ (primarily consisting of LBRs). Furthermore, we suspected that the effects of deleted genes encoding enzymes for the reactions within the SLBs would be more pronounced than the effect of genes encoding enzymes for uncoupled reactions. Indeed, this is confirmed by the analysis of the reaction type-based growth rates—the complex LBR (the number of element reactions of associated linear basis vector > 1), the simple LBR (the number of element reactions of associated linear basis vector = 1) and the uncoupled reactions (see Supplementary Information: Material S4)—and the average growth rates are 0.666, 0.773 and 0.813 h^−1^ (one-way ANOVA; *p* = 7.71E-28) (Fig 8e). Finally, we examined the cause for the observed differences among the flux variances, the number of simple cycles and the enzyme properties of LBRs. The flux variances are primarily explained by the multimeric enzymes and the topologically highly connected LBRs: SLBs are involved in a large number of simple cycles and few element reactions (Fisher’s exact test, *p* = 2.34E-21 and 0.0051, respectively) (Fig 8f). Therefore, the topological vulnerability of these reactions will result in functional variability.

## Discussion

We proposed Decrem by building the topologically decoupled metabolic network using SLB decomposition incorporating metabolic regulation by metabolites into GSMs, which approximate the kinetic fluxes of cell state-regulated reactions to constrain the feasible region of optimal flux distribution. Decrem effectively reduces the requirements for multi-omics data for genome-scale metabolic kinetic models. Compared to existing methods, Decrem demonstrates superior performance in predicting metabolic fluxes in three model organisms and growth rate in genome-scale knockout strains of *E. coli*. Therefore, it is an effective method for accurately depicting metabolic responses and exploring the self-adapting regulation mechanism of cellular perturbation.

By applying SLB decomposition, the (coupled) element reactions within identified SLBs display high coexpression among multiple growth conditions, indicating coordinated activation of topologically highly coupled reactions. Interestingly, similar approaches have been applied in identifying the non-redundant local functional units of metabolism, i.e., the minimal metabolic pathway or flux tope ^35,57^. A topological orthogonality principle was successfully used to design bioengineering strains with minimal interaction between desired product-associated pathways and metabolic components related to biomass synthesis ^58^. In addition, several specific topological constraints, such as removing the thermodynamically infeasible loops and decoupling two desired phenotypes, have been applied to GSMs to improve their metabolic production in recent studies ^59,60^. But Decrem is the first genome-scale topologically decoupled metabolic model for general applications, which clearly demonstrates how the topological preference of a metabolic network can guide the metabolic flux distribution.

To explore effective metabolic dynamic robustness or adaption to internal and external perturbations, metabolic kinetic models have attracted great attention by combining fluxomics, metabolomics and transcriptomics into a unified framework ^61-63^. However, construction of the GSM kinetic model is obstructed by the limited knowledge about kinetic parameters, e.g., *K*_m_, *K*_cat_, scarcity of metabolic regulators and paired genome-scale multi-omics data ^46,61^. To date, the largest metabolic kinetic model, k-ecoli457 of *E. coli*, contains only 457 reactions, 337 metabolites and 295 substrate-level regulatory interactions according to the computationally predicted kinetic parameters ^61^. Alternatively, by integrating the metabolite-TF regulatory regime, kinetic Decrem can predict growth rates and corresponding fluxes. A unique advantage of our kinetic Decrem is that it relies only on the experimental concentrations of identified essential metabolites, which serve as the key indicators of metabolic states and directly regulate enzyme activities or gene expression. Although some metabolites (e.g., those in glycolysis and the TCA cycle) have long been known to regulate enzyme activities in biochemistry textbooks, not much is known on the genome scale, even in model organisms. On the one hand, the regulatory metabolites of specific pathways in central metabolism prefer to frequently interact with the catalytic enzymes by activating or inactivating the functional domains to synchronously adjust the fluxes ^46,64^. On the other hand, several studies revealed that approximately 70% of the total variance in the promoter activity of metabolic genes in the central metabolism of *E. coli* can be explained by growth rate-derived global transcriptional regulation across multiple mutant strains ^48,65^. These findings suggest a potential relationship between the concentration of biomass-constituent metabolites and the expression of metabolic genes in the regulons of TFs. This relationship was verified by a recent study in which the identified metabolite concentrations were predicted by quantitative proteomics data ^66^. Overall, both the topology of metabolic networks and regulatory metabolites are utilized to identify coactivated or key regulated reactions, which produces a minimal set of regulatory constraints to develop the genome-scale kinetic models.

Compared to other methods for three model organisms, Decrem not only is the best in recovering the real flux distributions with high accuracy in a wide range of strains but also presents excellent predictions and explanations of the observed growth rate. This demonstrated the strong capability of Decrem to approximate the real intracellular state and to be used for designing high-yield mutant strains in bioengineering and synthetic biology. Decrem shows that there is a strong influence of metabolic network topology in the prediction of flux distributions; this phenomenon is found both in the reconstructed LBRs of Decrem and in the two versions of yeast metabolic models. A possible explanation for this observation is that the optimal flux distribution of a metabolic network is strongly determined by its topology, and the rewired or perturbed network structure will have different feasible regions. Moreover, Decrem could be applied to elucidate important regulatory branch points and the self-adaption of regulation mechanisms for the knockout strains, which is helpful for accurately predicting the potential target genes/reactions for designing bioengineering strains.

The main limitation of Decrem is that it requires a medium-scale set matched ^13^C-MFA flux paired with metabolite concentration data to construct reasonable kinetic models. Alternatively, the experimentally measured *K*_m_ and *K*_cat_ can also be used instead of the ^13^C-MFA flux. However, the well-constructed kinetic models are convenient for transfer to any other applications. Overall, Decrem, a topology and regulatory network-reinforced metabolic analysis model, can accurately predict phenotypes and uncover the complex regulation of cell metabolism.

## Methods

### Topology-decoupled reconstruction of the metabolic model

The original metabolic network consists of both coupled coactivated reaction cycles (e.g., TCA cycle) and simple linear chain reactions (e.g., biosynthetic reaction chains). We developed the model, Decrem, to capture the contribution of the coactivated and coupled reactions while preserving the consistency of the linear components. The detailed framework for our method is as follows(Supplementary Fig. 1):

#### Step 1: Identifying reaction cycle-based coupled substructures

Defining a bipartite graph derived from the metabolic network as *G*(*V*_*m*+*n*_, *E*), where the node set *V*_*m*+*n*_ includes both the *m* metabolite nodes and *n* reaction nodes, and the edge set *E* is composed of all the interactions between the metabolite and reaction nodes. We first built the similarity matrix *A*_*n*×*n*_ for *n* metabolic reactions based on the number of topological simple cycles of *G*. Specifically, the element *a*_*ij*_ of *A*_*n*×*n*_, which indicates the similarity between reactions i and *j* in the metabolic network, is measured by the number of simple directed cycles between the paired reaction nodes (*v*_*i*_, *v*_*j*_) of *G*. According to the similarity matrix *A*_*n*×*n*_ above, the BestWCut clustering algorithm ^34^ is then used to identify the dense substructures (also known as network communities) consisting of highly coupled reaction cycles. Here, the substructures are denoted as *C* = {*C*_*k*_|*k* = 1 … *K*}, where *C*_*k*_ is a subset of reaction index set [*n*] = 1, …, n, *k* is the index of substructures *C*_*k*_, and *K* is the total number of substructures. If we define *D*_*i*_ = ∑_*j*∈[*n*]_ *a*_*ij*_ as the weighted out-degree of reaction node *v*_*i*_, i ∈ [*n*], then the cluster degree *D*_*k*_ for subnetwork *C*_*k*_ could be defined as follows:

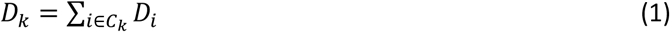

The generalized weighted cut (WCut) associated with *C* is obtained by minimizing *WCut*(*C*):

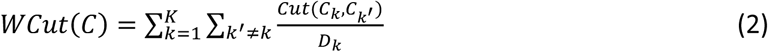

where

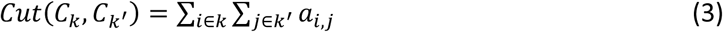

and *C*_*k*_′ is the complement set of the cluster *C*_*k*_

### Step 2: Reconstructing the decoupled representation of the identified substructure using a sparse linear basis

Inspired by the minimal metabolic pathways ^35^, we represented the highly coupled substructures with the SLBs of the null space of the corresponding stoichiometric matrix, which is biologically explained as the minimal and indecomposable coupled components, and satisfies the constraint of thermodynamic and mass balance of element reactions. Unlike infinite ordinary linear basis vectors of the null space of the stoichiometric matrix, there is a unique and globally optimal sparsest basis group of the null space ^35,67^. Briefly, the orthonormal null space 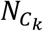 is initially defined by singular value decomposition (SVD) for the stoichiometric matrix 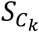 of the subnetwork *C*_*k*_. Here, additional artificial exchange reactions are introduced in 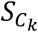 to maintain the mass balance of reactions in the subnetwork *C*_*k*_ (more details can be found in Supplementary Information: Method S1). Then, the column vectors of the orthonormal null space 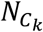 are iteratively replaced by the minimal element reactions that span the removed subspace of vectors. This process is repeated until all the nonzero entries in 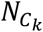 converged on a minimum ^35^. Here, we utilized the advantage of sparse regularization of the null space 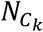 to solve the minimum L1-norm of the null space 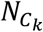 of 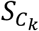 ^67^. The detailed process is showcased in Supplementary Information: Method S1, in which 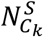 is a minimal sparse basis representation of 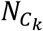 in at most 2*r*_*k*_ linear programming optimization runs (where 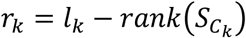, *l*_*k*_ is the number of columns (reactions) in 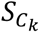). If we assume *x* ∈ R^*n*^ and then each linear programming problem can be formulated as follows:

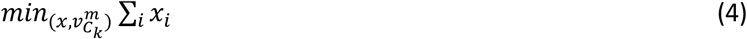

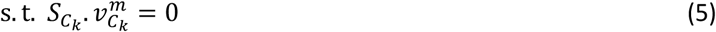

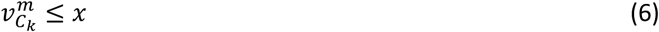

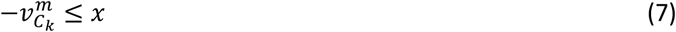

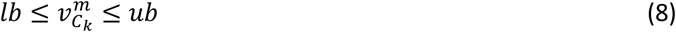

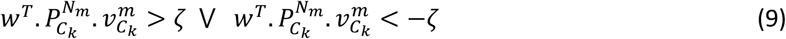

where 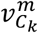 is the SLB of the null space of 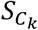 at the *m*^th^ running, and *lb* and *ub* are the lower and upper bounds of 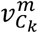, respectively. The constraint of formula (9) ensures that 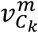 is linearly independent of the previous *m* − 1 SLBs, and w represents a vector of random weights. Here, we employed uniform random weights, and ζ is a small positive constant, e.g., 1.0^−3^. 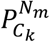 is a projection matrix onto the sparse null space 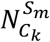, and 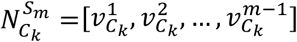. More details of this process are provided in Supplementary Information: Method S1.

The SLBs of 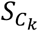, assumed as 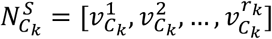, can be solved by the above algorithm. Then, the highly coupled subnetwork *C*_*k*_ can be substituted to construct the mutually independent “linear basis reactions” with the following formula:

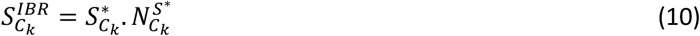

where 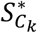 and 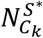 are derived from 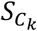 and 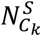 after removing the artificial exchange reactions, respectively, and *IBR* indicates the reconstructed independent “linear basis reactions”.

### Step 3: Establishing prediction for the decoupled metabolic model

In this step, we reformulated FBA to adapt the reconstructed decoupled metabolic network *S*^*IR*^. The key problem is to determine the flux bounds for each “linear basis reaction”:

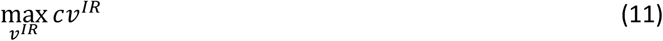

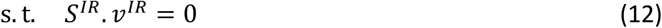

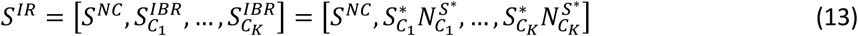

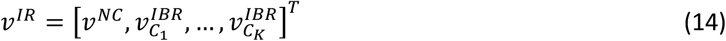

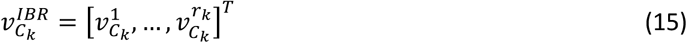

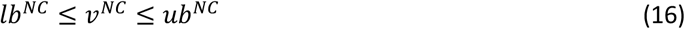

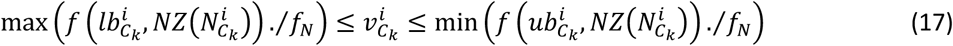

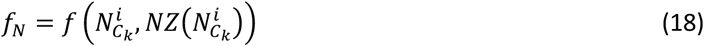

where *c* and *v*^*IR*^ represent the objective function and optimal metabolic flux of *S*^*IR*^, respectively. The superscript *IR* represents the linearly independent reaction-derived metabolic network, and *NC* represents the uncoupled reactions (which are composed of linear reaction chains) of the original metabolic network. *r*_*k*_ is the number of columns of 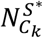, and *K* is the total number of highly coupled reaction subnetworks identified in *step* 1. 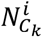 is the ith column of 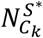, and 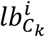 and 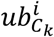 are the lower and upper bounds of the reaction indicated by 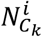, respectively. 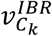 is the flux vector of all the “linear basis reactions” of subnetwork *C*_*k*_, and 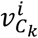 is the ith flux of 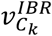. Among them, *i* ranges from 1 to *r*_*k*_, and k ranges from 1 to *K*. Then, the fluxes of reactions in the original metabolic network (element reactions) of linear basis vectors will be recovered by the formula 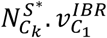 according the optimal solution of Decrem outlined above.

The function *NZ*(.) takes the index of nonzero elements of the input vector, and the function *f*(*v, I*) takes elements indexed by the input indicator *I* from the input vector *v*. Therefore, 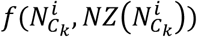 represents the nonzero partition coefficient of each element reaction composed of the *i*^th^ SLB of subnetwork *C*_*k*_, which is indicated by the nonzero terms of 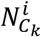. The formula 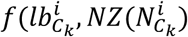 represents the lower bounds of the element reactions composed of the *i*^th^ SLB of the subnetwork *C*_*k*_, and 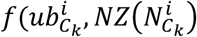 M represents the upper bounds. In summary, Decrem forces the metabolic fluxes of highly coupled reactions to be incorporated in optimization by representing them as independent linear basis vectors.

### Gene expression estimation

Forty-five candidate global and local regulatory metabolites of *E. coli* are collected through KEGG pathway analysis and a literature survey ^50,51,68^. These candidates are then categorized into two clusters by hierarchical clustering analysis over the expression profile of their concentrations across 24 mutant strains. Then, the possible regulatory relationship between two identified metabolite groups and the expression of 85 genes are inferred by PLSR ^69^, which selects the nonredundant and independent factors to maximize the correlation of response variables, using stepwise principal component regression. Furthermore, PLS is used to discover the fundamental quantitative relations between two (expression) observation variable sets, and the general underlying model of multivariate PLS is as follows:

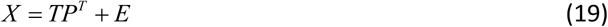

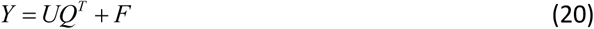

where *X* is an *n* × *m* matrix of predictors (each group metabolites concentrations), and *Y* is an *n* × *p* matrix of responses (genes expression); *T* and *U* are *n* × *l* matrices and projections of *X* (the *X* score, component or factor matrix) and projections of *Y* (the *Y* scores), respectively. *P* and *Q* are *m* × *l* and *p* × *l* orthogonal loading matrices, respectively. Matrices *E* and *F* are the error terms, assumed to be independent and identically distributed random normal variables. The decomposition of *X* and *Y* was performed to maximize the covariance between *T* and *U*.

Finally, the significant metabolite profile and corresponding explicable gene-metabolite regulatory relationships are filtered by setting the proper correlation threshold and the statistical test is built based on random sampling (see Supplementary Information: Material S3).

### The metabolite concentration-derived linearized kinetic model

#### Step 1: The metabolic kinetic model

In this section, we derived a complete reversible rate law for arbitrary reactant stoichiometries. When considering the constraint of thermodynamics and metabolite regulation ^46,70^, we can rewrite the Michaelis–Menten kinetics for a reversible reaction S <=> P as follows:

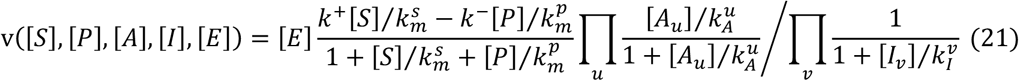

where [*E*] is the concentration of enzyme active sites, [*S*] and [*P*] are the concentrations of substrates and products, 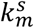 and 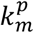 are the affinities of the reactants for this enzyme, *k*^+^ and *k*^−^ are the maximal forward and reverse catalytic rate constants, [*A*] and[*I*] are the concentrations of activators and inhibitors, and 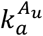 and 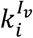 are their corresponding affinities. The positive and negative terms in the numerator are associated with the forward and backward rates, respectively.

We next applied the metabolic thermodynamics constraints given by the Haldane relationship to simplify the term for the backward rate:

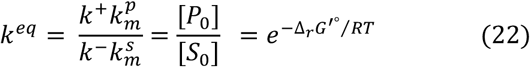

where 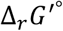 is the standard Gibbs energy of the reaction (and does not depend on the enzyme parameters). Using this equality with the above rate law, we can obtain the following:

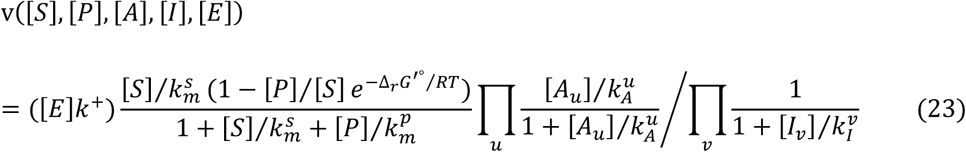

### Step 2: The optimal strategy of the linearized kinetic model

In this section, we sought the simplified representation of equation (23) based only on the associated metabolite concentrations. First, we took the negative logarithmic operation of both sides of equation (23) and reorganized the right-hand terms:

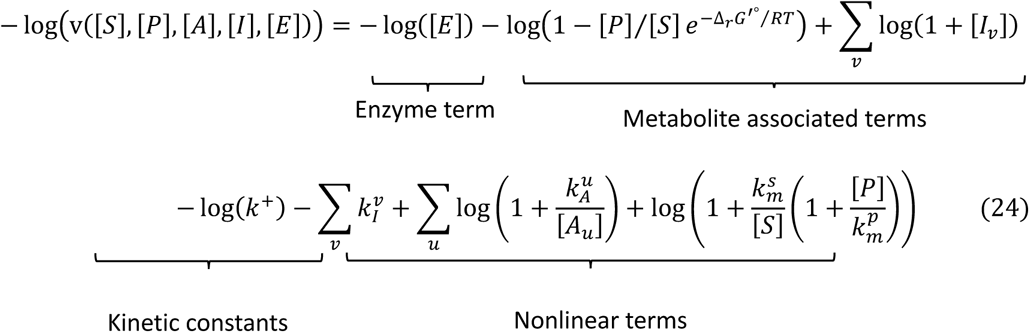

This model can be solved by collecting the corresponding kinetic parameters, enzyme expression and metabolite concentrations; however, those matched data are often not available in practice, and the metabolic regulators are often unknown. An alternative is to approximate the optimal parameters according to the machine learning method. Specifically, the cell growth state-regulated enzyme expression can be represented as the linear combination of the concentrations of biomass composition (see Supplementary Information: Material S3), which can be marked as 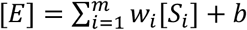, where *S*_*i*_ is the concentration of the *i*th indicator metabolite of the biomass identified in the “Gene expression estimation” section. In addition, we reexamined the nonlinear terms of equation (24) based on the knowledge that systemic experimental analysis revealed that [*S*] was ≥ 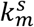 for almost all of the metabolites in the central metabolism of three model organisms ^47^; hence, we have 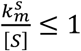.

Situation 1: For the growth state-regulated irreversible reaction, if we have 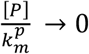, and 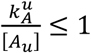, then the Taylor expansion of equation (24) is as follows:

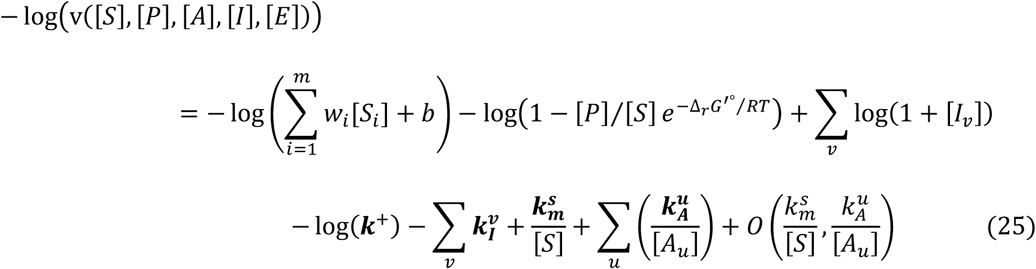

Situation 2: For the generalized reversible reaction in which the enzyme is substrate or product saturated (or both): 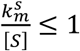 and 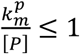. Assuming *v* > 0, we have the following:

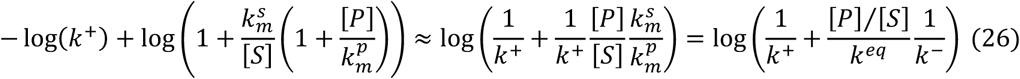

Here, [*P*]/[*S*] < *k*^*eq*^ according to the reaction Gibbs energy, 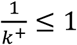 and 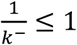 for the significantly regulated reactions, and 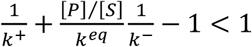; so, the Taylor expansion of(26) is as follows:

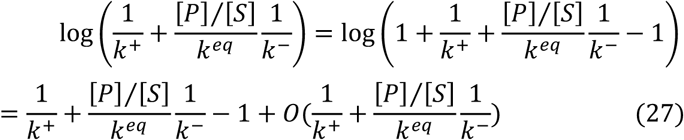

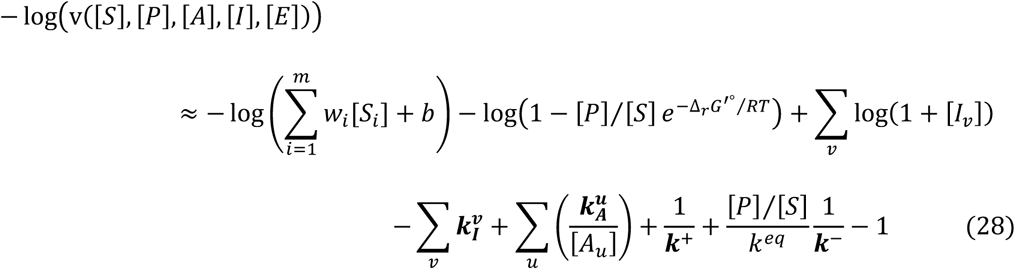

where all the bold variables in equations (25) and (28) are kinetic parameters. Similarly, when *v* > 0, we have 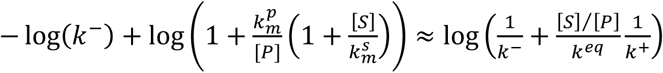. This result can be expanded to multisubstrate/multiproduct reactions.

Overall, when we neglected the infinitesimal of higher order, the identified regulators and optimal kinetic parameters in models (25) and (28) can be solved with a Bayesian linear regression-derived model selection method, i.e., the SIMMER package ^46^, which first builds the regulator-free kinetic model, and then uses the BIC criteria to identify the real regulators from the candidate metabolite set (PG metabolites) by evaluating the additional ability to account for the empirical fluxes when integrating a new candidate regulator into the kinetic model in an incremental way (see Supplementary Information: Methods S3). Subsequently, the optimal model is expanded to any other application.

### Step 3: The kinetic Decrem

Finally, the parameterized kinetic model is utilized to describe the growth-associated key regulated reactions in central metabolism. The genome-scale flux distribution is predicted by the kinetic regulated flux-constrained Decrem method, i.e.,

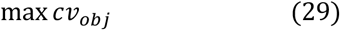

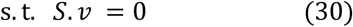

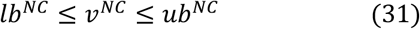

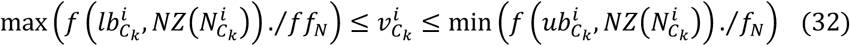

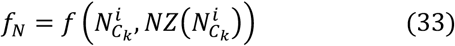

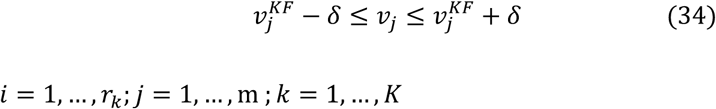

where 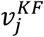 is the *j*-th kinetic flux of the *m* key regulation reactions, and *δ* is the tolerance of kinetic fluxes. The 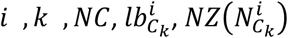 M values can be found in *step* 3 of the section “Topology-decoupled reconstruction of the metabolic model”. Among them, *i* ranges from 1 to *r*_*k*_, *j* ranges from 1 to m, and k ranges from 1 to *K*.

## Data Availability

All data used are publicly available. The original and reconstructed metabolic models are available online: original metabolic models are available at BIGG models http://bigg.ucsd.edu/models, and reconstructed metabolic models of the three reconstructed models, iAF1260, iMM904, iDN750, are available at https://github.com/lgyzngc/Decrem-1.0.git. All used exchange reactions, nutrient uptake, experimental growth rates, 13C fluxes and gene expression for Decrem modeling and metabolic simulation are found in supplementary tables. And the LS-MS data is sourced from https://www.ebi.ac.uk/biostudies for genome-scale mutant strains of E. coli.

## Code Availability

Decrem is implemented as a Matlab package, the source code, user tutorial and demo are available at: https://github.com/lgyzngc/Decrem-1.0.git.

## Acknowledgements

The authors thank Schellenberger J’s open-source Matlab package: Cobra, and Saa PA’s open-source Matlab package: Fast_SNP for the integration of Decrem. This work was supported by the National Natural Science Foundation of China (61872418) and Natural Science Foundation of Jilin Province (20180101050JC).

## Materials & Correspondence

Correspondence and requests for materials should be addressed to G.L., W.D. or H.C.

## Supplementary information

**Supplementary Fig. 1:** The workflow of Decrem; **Supplementary Fig. 2**: The accumulation flux distribution of according the Yeast iMM904 model (red) and reconstructed decoupled model (blue) by the flux variability analysis; **Supplementary Fig. 3**: Comparison of activated reactions of TCA cycle among 13C MFA (A), Decrem (B) and FBA (C). Solid line indicate the activated reactions and dash line indicate the reactions having a zero flux; **Supplementary Fig. 4**: The workflow of kinetic regulation model of Decrem; **Supplementary Fig. 5**: The concentration profiles of 45 candidate regulating metabolites according to hierarchical clustering, under 24 mutant strains. The red and blue rectangles mark the identified two high-correlation groups; **Supplementary Fig. 6**: The consistency of 32 genes between measured and predicted expression levels by the PLSR methods; **Supplementary Fig. 7:** The growth rate distribution of the 1030 metabolic genes knockout strains predicted by the conventional methods: FBA, MOMA and Decrem, respectively, without any external flux constraints; **Material S1**: The data processing in the section of “Prediction of metabolic fluxes in response to environmental changes”; **Material S2**: The data processing in the section of “Prediction of metabolic fluxes and growth rates in yeast knockout strains”; **Material S3**: The workflow and data processing in the section of “Building a metabolic kinetic model incorporating metabolite-TF regulation”; **Material S4**: The workflow and data processing in the section of “Growth rate estimation for E. coli genome-scale deletion strains with kinetic Decrem”; **Method S1**: The Decrem algorithm; **Method S2**: The Parsimonious Decrem; **Method S3**: Identifying the high confidence regulators by Bayesian model section method.

**Supplementary Table 1**. The annotation of reconstruction decoupled metabolic models, reconstructed models can be found at Data availability.

**Supplementary Table 2**. Nutrient uptake reactions for three microorganisms under six growth conditions.

**Supplementary Table 3**. The 13C MFA fluxes for three microorganisms under six growth conditions.

**Supplementary Table 4**. The dataset used to predict the fluxes and growth rates for the Yeast 38 deletion strains.

**Supplementary Table 5**. The correlation statistic between the experimental and predicted metabolic fluxes and growth rates for the Yeast 38 deletion strains.

**Supplementary Table 6**. The dataset used to build and validate the hierarchical regulation model.

**Supplementary Table 7**. The dataset used to construct kinetic Decrem to predict the growth rates on genome-scale deletion strains.

